# Evolutionary trajectories of cohesin gatekeepers across the Tree of Life

**DOI:** 10.1101/2025.05.08.652912

**Authors:** Daniela Quiroz, Alice Pierce, Grey Monroe

## Abstract

Eukaryotic genomes are physically organized into chromatin, which regulates virtually every dimension of genome biology, including replication, recombination, transcription, and DNA repair. Precocious Dissociation of Sisters 5 (PDS5) protein is a key regulator of the cohesin complex, responsible for generating higher-order chromatin structures within and between chromosomes. In yeast and vertebrates, PDS5 can lead to cohesin establishment or disassembly, depending on its interaction with other proteins. In this study, we explored the diversity of PDS5 proteins across eukaryotes; HEAT-repeats forming a hook-like structure (PDS5 domain), followed by a predicted disordered region of variable lengths, are observed across the Tree of Life. Interestingly, two homolog types (A and B) with signs of varying chromatin localization emerged independently in later branches of animal and plant kingdoms, in vertebrates and vascular plants, respectively (∼400-500 MYA). We also investigated the emergence of two plant-specific features in the PDS5 proteins: 1.) An ancestral acquisition of a Tudor domain with conserved H3K4me1-binding amino acids in land plants 2.) A truncated version of PDS5 proteins that lacks essential features of PDS5 domain and is involved in the RNA-directed DNA methylation (RdDM) plant-specific pathway which emerged in seed plants. This work provides the most comprehensive study of PDS5 diversity, and suggests the emergence of specialized cohesin regulatory mechanisms that could shape distinct strategies for genome organization across eukaryotes.

**Short summary:** Here, we analyzed the diversification of PDS5 proteins; sequence, domain content, and structure, providing new observations that evidence different evolutionary trajectories of this protein across the Tree of Life. These results suggest neofunctional evolution in cohesin regulatory proteins that could shape distinct strategies for genome organization across eukaryotes.

## Introduction

Organizing large genomes within the nucleus is a highly regulated process essential for numerous cellular functions, including gene expression, DNA replication, repair, recombination, and chromosome segregation. One key element is the **cohesin complex**, a multi-protein structure involved in sister chromatid cohesion (SCC) and higher-order folding of chromatin, which helps organize the genome into topologically associated domains (TADs) and territories (Bolaños-Villegas et al., 2017; Choudhary & Kupiec, 2023; Rittenhouse & Dowen, 2024; Yu et al., 2022). The cohesin complex, conserved across eukaryotes, includes members of the Structural Maintenance of Chromosomes (SMC) family, consisting of two SMC proteins, SMC1 and SMC3, which form a V-shaped heterodimer, and a kleisin subunit SCC1 (RAD21 in most eukaryotes, SYN proteins in *Arabidopsis thaliana*), which bridges the two arms of the V, creating a ring-like structure capable of entrapping DNA. In addition, cohesin can interact with several regulatory subunits, that control the establishment and disassembly of the complex, such as SCC3 (STAG1 or STAG2 in mammals), SCC2 (NIPBL in vertebrates), SCC4 (MAU2 in vertebrates) and **PDS5** (alternatively SPO76 or RDM15), also known as the **cohesin gatekeeper**. (Bolaños-Villegas et al., 2017; Choudhary & Kupiec, 2023; Rittenhouse & Dowen, 2024; Yu et al., 2022).

PDS5 is a large protein composed primarily of HEAT-repeats, formed by two alpha helices connected by a short loop, that fold into a superhelical structure that facilitates protein–protein interactions (Yoshimura & Hirano, 2016). It stabilizes cohesin by interacting with SCC1, while its association with WAPL promotes cohesin disassembly by opening the SMC3-SCC1 ring. After DNA replication, cohesin is further stabilized through SMC3 acetylation, a process indirectly promoted by PDS5 via recruitment of CDCA5 (Sororin), which displaces WAPL (Morales et al., 2020; Nishiyama et al., 2010; Wutz et al., 2017).

SCC1 and WAPL are the only proteins interacting with PDS5 for which experimental structures are available (Lee et al., 2016; Ouyang et al., 2016). The SCC1-PDS5 complex (PDB: 5F0N, 5F0O) is based on yeast sequences, whereas the WAPL-PDS5 interaction (PDB: 5HDT) is derived from human sequences. Although no additional direct structural evidence exists for PDS5 interactions, indirect evidence suggests that PDS5 associates with numerous proteins both within and outside the cohesin complex across eukaryotes (Supplementary Table 1).

In budding yeast, knocking out PDS5 results in severe defects in sister chromatid cohesion, leading to premature chromatid separation, mitotic arrest, and ultimately cell lethality (Hartman et al., 2000; Stead et al., 2003). Other experiments suggest that PDS5 has a role in coordinating DNA replication and sister chromatid cohesion establishment (Choudhary et al., 2022; Choudhary & Kupiec, 2023) In vertebrates, there are two PDS5 paralogs, PDS5A and PDS5B. Mice lacking either PDS5 homolog have a lethal embryogenic phenotype (Carretero et al., 2013). In Xenopus egg extracts, depletion of PDS5A and PDS5B results in mitotic chromosomes that display normal arm cohesion but weakened centromeric cohesion, accompanied by an abnormal retention of cohesin (Losada et al., 2005). Depletion of these proteins also has a weakening effect on chromatin loops in human cells (Yu et al., 2022). While both homologs contribute to telomere and arm cohesion, PDS5B is essential and specific for centromere cohesion. (Carretero et al., 2013; Zhang et al., 2021). Besides those distinctions, PDS5A and PDS5B differentially affect gene expression without altering cohesin localization (Arruda et al., 2022).

The model plant *Arabidopsis thaliana* (Arabidopsis, hereafter) encodes for five homologs of PDS5 (A to E), which have redundant and distinct functions. Depletion of these PDS5 proteins leads to severe effects on development, fertility, somatic homologous recombination (HR), and DNA repair (Bolaños-Villegas et al., 2017; Pradillo et al., 2015). Newly emerged evidence demonstrates that Arabidopsis PDS5 proteins play a negative role in TAD-like domain formation (Göbel et al., 2024). Interestingly, the PDS5C and PDS5E homologs have been associated with the plant-specific RdDM pathway, and PDS5C Tudor domain has been shown to bind H3K4me1 (Niu et al., 2021; Takei et al., 2024).

Emerging evidence shows surprising diversity in PDS5 proteins, such as acquiring a reader domain in PDS5 plant homologs. This evidence points to lineage-specific adaptations and raises questions about the evolutionary origins and diversification of PDS5 proteins’ domain architecture and functions. In this study, we will investigate PDS5 protein diversity across the eukaryotic Tree of Life. In particular, we will address domain content, secondary structure differences, and amino-acid diversity of the cohesin gatekeeper.

## Results & Discussion

### Arabidopsis PDS5 homologs have Tudor domains. Homologs C, D, and E exhibit shorter HEAT-repeat regions

A comparative analysis of PDS5 orthologs in *Saccharomyces cerevisiae, Arabidopsis thaliana*, and *Homo sapiens* reveals distinctive domain compositions and structural features (Fig. 1). In yeast, a single PDS5 homolog is present, characterized by an HEAT repeat-rich PDS5 domain (predicted pfam accession PF20168), forming a hook-like structure. In humans, two homologs, PDS5A and PDS5B, share similar HEAT repeat-rich PDS5 domains. However, they differ from the yeast homolog and from each other in the length and content of their intrinsically disordered regions (IDRs), with PDS5B exhibiting longer IDRs. Zhang et al. (2021) predicted multiple features within the C-terminal IDRs of vertebrate PDS5A and PDS5B homologs, including nuclear localization signals, numerous phosphorylation sites, and, in the case of PDS5B, two AT-hook motifs with potential to bind AT-rich DNA. Despite an overall sequence identity of ∼70% between PDS5A and PDS5B, these differences in IDR length and motif content may underlie their distinct chromatin localization and interaction profiles. IDRs are known to play crucial roles in transcriptional regulation by mediating subnuclear localization, dynamic protein-protein interactions, and assembly of large complexes (Cermakova & Hodges, 2023). In nuclear proteins, IDRs often contribute to chromatin organization, transcriptional control, and formation of dynamic condensates via liquid–liquid phase separation (Brodsky et al., 2020; Homma et al., 2012; Miao & Chong, 2025). Although no experimental data yet directly link these IDR features to PDS5 localization and function, their conservation and complexity highlight the need for future experimental investigation (Arruda et al., 2022; Carretero et al., 2013; Zhang et al., 2021).

**Figure 1.**
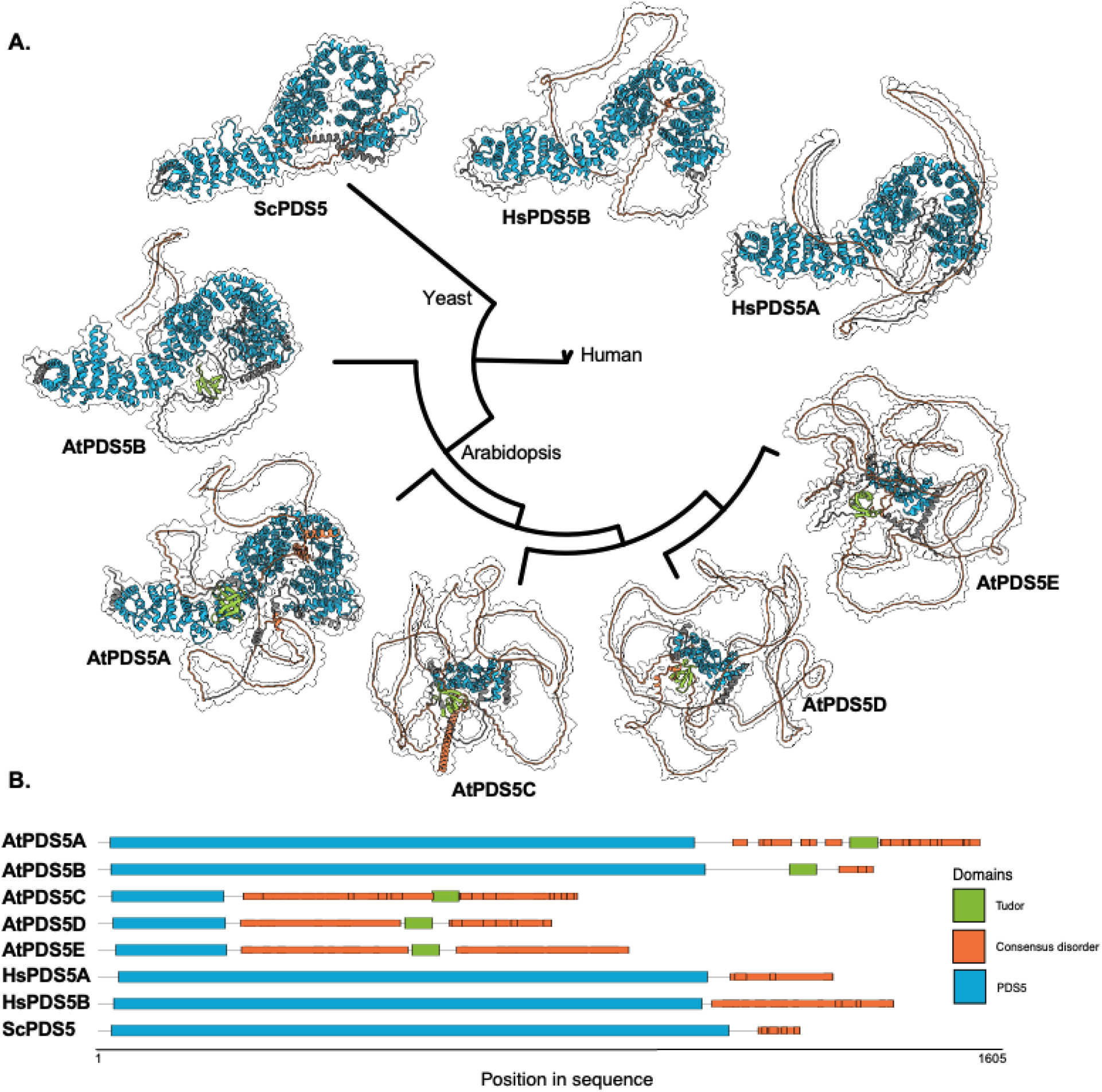
Phylogenetic and Structural Analysis of PDS5 Proteins Across Yeast, Human, and *Arabidopsis thaliana*. **A**. Maximum-likelihood phylogenetic tree of representative PDS5 proteins, including *Saccharomyces cerevisiae* (ScPDS5), *Homo sapiens* (HsPDS5A and HsPDS5B), and *Arabidopsis thaliana* (AíPDS5A, AtPDS5B, AíPDS5C, AíPDS5D, AtPDS5E). The tree was constructed using IQ-TREE and the best-fit model (*Q*.*insect*+*F*+*R10*), rooted in yeast (*ScPDS5*). The structures correspond to AlphaFold3 predictions of each protein, visualized with their respective Tudor domains (green), consensus disordered regions (orange) and PDS5 domains (blue). **B**. Domain architecture of PDS5 proteins based on InterProScan results. The domains are highlighted: Tudor domains (green), consensus disordered regions (orange) and PDS5 domains (blue). Positions correspond to sequence coordinates from 1 to 1605 amino acids.

In *Arabidopsis thaliana*, five PDS5 homologs (PDS5A to PDS5E) have been identified (Pradillo et al., 2015). PDS5A and PDS5B possess HEAT repeat-rich PDS5 domains comparable in size to those in yeast and humans (900-1100 aa), while PDS5C to PDS5E exhibit smaller PDS5 domains (200 - 300 aa). This size variation suggests functional divergence, with PDS5C-E potentially playing roles in pathways unique to plants, such as RNA-directed DNA methylation (RdDM) (Niu et al., 2021; Takei et al., 2024), although new evidence suggests that these smaller PDS5 proteins also play a role in cohesin (Göbel et al., 2024); however, the specific mechanism remains unclear, as these proteins lack several of the structural features traditionally described for canonical PDS5 proteins. Additionally, all *Arabidopsis* PDS5 homologs contain IDRs of variable sizes and a Tudor reader domain, the latter shown to interact with H3K4me1 in PDS5C homolog (Niu et al., 2021).

Given the role of cohesin in homology-directed repair of double-strand breaks, the Tudor-H3K4me1 of PDS5 may contribute to lower mutation rates in gene bodies by promoting epigenome-directed DNA repair (Quiroz et al., 2024). These features highlight unique and potentially adaptive functions of PDS5 proteins in plants.

### Conserved Single-Copy of PDS5 Across Eukaryotes, with Lineage-Specific Duplications in Vertebrates and Vascular Plants

We developed a computational pipeline to identify and curate PDS5 homologs across eukaryotes (Supplementary Fig. 1A). Using BLASTp with yeast, human, and Arabidopsis PDS5 queries, we identified 24,329 putative homologs across 7,918 eukaryotic genomes. After filtering by domain content (Pfam PF20168), sequence redundancy (>85% identity within species), and domain size distribution per major clade, 6,165 high-confidence PDS5 sequences from 4,020 species were retained for phylogenetic analysis. This represents the most complete analysis of PDS5 diversity conducted to date, complementing previous, smaller-scale analyses (Takei et al., 2024; Zhang et al., 2021).

We analyzed the distribution of PDS5 domain sizes across major eukaryotic groups (Supplementary Fig. 1B). Protists and Fungi have variable domain sizes (1,030–1,200 amino acids), while Animals have a highly conserved domain size (1,030–1,050 amino acids). In Plantae, we observed a bimodal distribution, with peaks around 1,000 and 230 amino acids, consistent with the sizes of the previously described Arabidopsis PDS5 homologs. Based on these patterns, group-specific size thresholds were applied to filter out likely truncated sequences, ensuring consistency for downstream evolutionary analyses (see Materials and Methods). While we acknowledge that strict size-based filtering may exclude true PDS5 homologs with atypical domain sizes, we prioritized the retention of sequences with conserved structural features to maximize reliability in evolutionary inferences. This is especially important in order to avoid artifacts introduced by genome annotation errors.

We then constructed a maximum likelihood phylogenetic tree to assess the evolutionary relationships of PDS5 homologs across eukaryotes. The tree revealed PDS5 as a conserved protein across diverse eukaryotic lineages, including basal clades such as protists, basal fungi, opisthokonta, basal metazoans, and rhodophytes, as well as derived groups like vertebrates and vascular plants, reflecting previously described essential role of PDS5 among eukaryotes (Carretero et al., 2013; Hartman et al., 2000; Morales et al., 2020; Pradillo et al., 2015; Yu et al., 2022). The phylogenetic tree, constructed using the PDS5 domain from up to three representative species per class, showed the expected clustering of major eukaryotic groups (Fig. 2A). Protists (e.g., Stramenopiles) occupied the most basal branches, with other basal lineages clustering outside the main eukaryotic clades (Fungi, Animalia, and Plantae) reflecting the deep evolutionary history of PDS5 across eukaryotes. Phylogenetic analysis also showed that most eukaryotic clades retain a single PDS5 homolog, consistent with a single PDS5 encoding gene in the ancestor of eukaryotes (Fig. 2B). However, conserved duplication events were identified in two complex-eukaryotic major clades: vertebrates and vascular plants. Our analysis likely remains unaffected by recent whole-genome duplications (e.g., in polyploid plants), as we only counted PDS5 homologs that display distinct sequences (e.g., >15% sequence difference), rather than near-identical copies arising from recent duplications.

**Figure 2.**
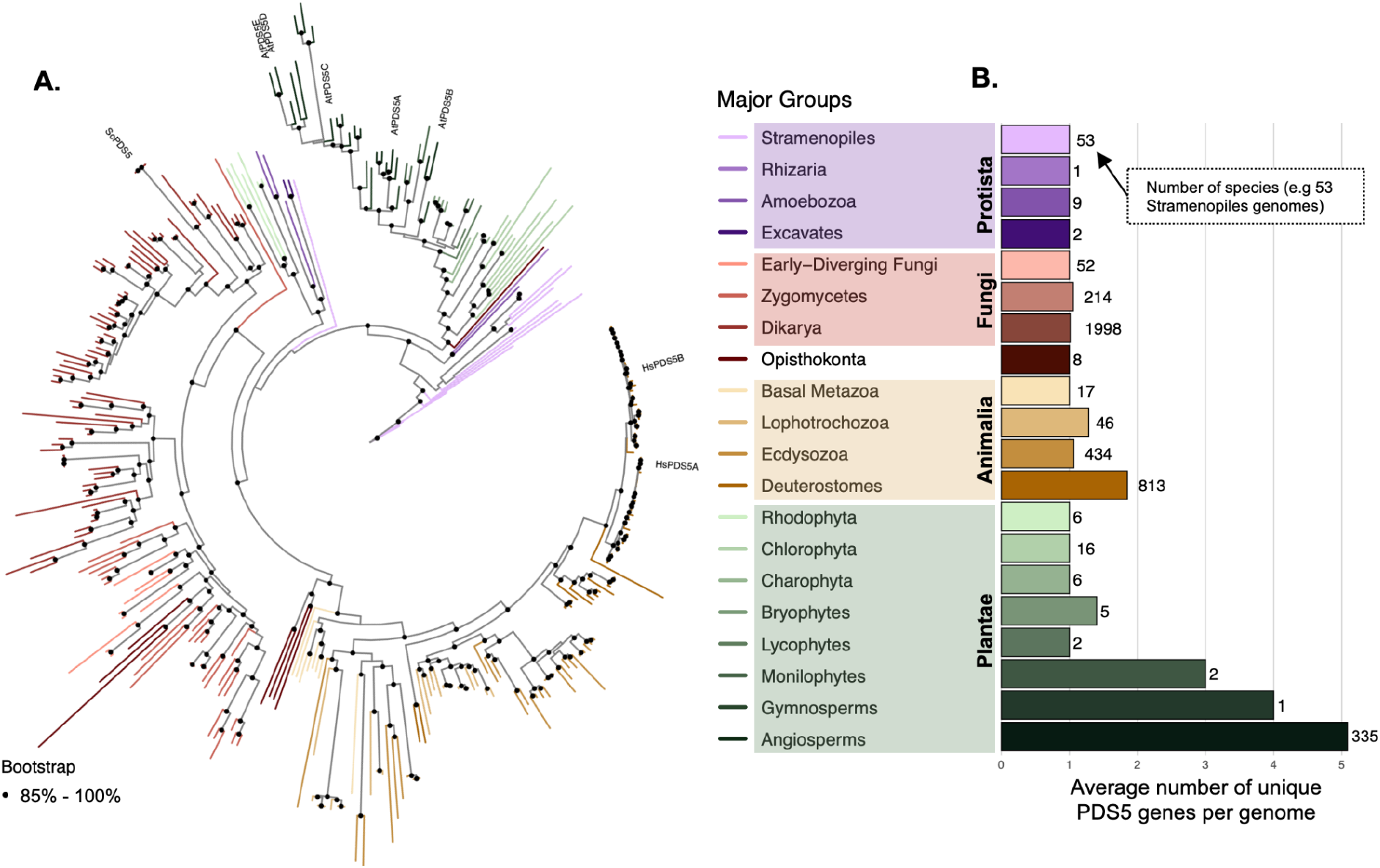
Maximum Likelihood Tree of PDS5 Proteins Across the Tree of Life. **A**. Maximum-likelihood phylogenetic tree of the PDS5 protein family constructed using the best-fit evolutionary model (*Q*.*insect*+*F*+*R10*). The tree is rooted in Stramenopiles and includes 1–3 representative species per class(128 classes in total). Black dots at the nodes represent bootstrap support values higher than 0.85 (based on 1,000 replicates). **B**. Average counts of PDS5 proteins per species across Major Groups within the Tree of Life. Each bar represents the mean protein count for species in each clade, with the number of species shown at the end of the bar.

In vertebrates, PDS5A and PDS5B homologs emerged early, with representatives already present in jawless vertebrates such as *Petromyzon marinus* (Supplementary Fig. 5), and exhibit high sequence conservation across species (Supplementary Fig. 6). As mentioned previously, sublocalization of vertebrate homologs PDS5A and PDS5B has been shown to target different chromosome structures, with PDS5B being more exclusively associated with centromeric regions during mitosis (Carretero et al., 2013; Yu et al., 2022). Further subfunctionalization of these two homologs is evident in their distinct influence on gene expression in mouse embryonic cells (Arruda et al., 2022). The presence of both homologs being highly conserved across vertebrates reflects a crucial subfunctionalization or neofunctionalization of the emergent homolog.

Similarly, two homologs, which have also been named previously PDS5A and PDS5B arose prior to the divergence of ferns in vascular plants. In the case of plants homologs A and B, no experimental evidence has shown distinct subcellular localization or clear signs of neofunctionalization to date. However, indirect inference suggests functional differentiation based on the differing effects of PDS5A and PDS5B knockouts in the TAD-like domains in Arabidopsis. Where the presence of PDS5A, but not PDS5B alone, leads to the genome-wide suppression of TAD-like structures. (Göbel et al., 2024). Furthermore, co-immunoprecipitation experiments in Arabidopsis find that PDS5A, but not PDS5B, is found in association with PDS5C; therefore, PDS5A and PDS5B could have a distinct subcellular localization (Göbel et al., 2024; Takei et al., 2024).

Another homolog type, “PDS5C class”, emerged during the diversification of spermatophytes (seed plants, including gymnosperms and angiosperms), with additional duplications observed in specific lineages within flowering plants. These smaller homologs have also been associated with the cohesin complex (Göbel et al., 2024; Pradillo et al., 2015; Takei et al., 2024). However, the precise mechanism of action for these truncated PDS5 homologs remains unclear, as the canonical hook-like structure that mediates cohesin binding is absent, and the HEAT-repeat interacting surface is significantly reduced. Interestingly, PDS5C and related truncated homologs have also been implicated in plant-specific RNA-directed DNA methylation (RdDM) pathways. Niu et al. (2021) demonstrated that PDS5C interacts with NRPE3B, a Pol V-specific subunit, and that it is essential for siRNA accumulation and DNA methylation at a subset of RdDM loci, establishing a direct connection between histone marks and Pol V activity. Similarly, Takei et al. (2024) identified PDS5C and PDS5E homologs in Arabidopsis as angiosperm-specific that interact with AGO1 and are essential for 24-nt siRNA production, to establish RdDM at transposable elements (TEs). Loss of PDS5C/E function leads to defects in siRNA accumulation and CHH DNA methylation, suggesting a unique regulatory role distinct from their canonical cohesin-associated functions. These findings suggest a new role of PDS5C/E, which could reflect an evolutionary adaptation to manage transposable element (TE) expansion in angiosperms (Takei et al., 2024). Beyond plants, evidence suggests a broader interplay between the cohesin complex and non-coding RNAs across eukaryotes. Kuru-Schors et al. (2021) reviewed multiple examples where small RNAs (sRNAs) and long non-coding RNAs (lncRNAs) regulate cohesin components, influencing gene expression, genome stability and chromatin dynamics (Kuru-Schors et al., 2021). Future work could explore whether specific small RNA subclasses contribute directly to cohesin establishment or stabilization in plants, and clarify a potential interplay between RdDM pathway and cohesin dynamics.

### Emergence of Tudor Domain and PDS5C Homologs Reflects a Plant-Specific Diversification of PDS5

Domain content analysis across all identified PDS5 homologs revealed a deeply conserved architecture with PDS5 domain (PF20168), and intrinsically disordered regions (IDRs). Some clades also present Tudor domains across lineages (Supplementary Fig. 2). No additional conserved domains were identified beyond these three features.

In protists and basal fungi, PDS5 homologs showed extensive variability in both IDR content and PDS5 domain size (Supplementary Figs. 3 and 4), suggesting diversification in cohesin organization and regulation in early eukaryotes. While more recently diverged fungi showed variability in PDS5 sequence (Supplementary Fig. 6), and IDR length (Supplementary Fig. 4), suggesting an adaptive divergence of both regions, suggesting potential lineage-specific adaptations, although experimental evidence will be required to confirm their functional relevance.

In animals, PDS5 homologs were highly conserved across major clades, with vertebrates displaying two paralogs (PDS5A and PDS5B) that arose early in vertebrate evolution (e.g., Petromyzon marinus; Supplementary Fig. 5). Both paralogs exhibit minimal sequence divergence (Supplementary Fig. 6), reflecting essential conserved roles in cohesin regulation in the PDS5 domain as previously evidenced in several vertebrate organisms (Arruda et al., 2022; Morales et al., 2020; Yu et al., 2022; Zhang et al., 2021).

In plants, we find that the most parsimonious model involves three key transitions in the PDS5 evolutionary trajectory (Fig. 3A). **First** (Figure 3, Step 1), a Tudor domain likely emerged in the common ancestor of all land plants, as. we found it within the basal Charophyte clade (*Closterium, Klebsormidium, Chara* species) (Fig. 3A–B; Supplementary Fig. 2A). This acquisition likely occurred between 500–700 million years ago (MYA), prior to the diversification of vascular plants, and represents a major evolutionary innovation linking PDS5 to histone-reader function (Quiroz et al., 2024). To explore potential sources of the PDS5 Tudor domain, we examined other genes in the Arabidopsis genome encoding Tudor domains and found that the plant-specific Tudor domain of the mismatch repair protein MSH6 shares considerable similarity. Interestingly, recent work (Monroe et al., 2025) suggests that the Tudor domain in MSH6, likely originated in the common ancestor of Viridiplantae, cryptomonads, and SAR lineages, thus preceding the putative PDS5C Tudor acquisition event we find here. This raises the possibility that the PDS5 Tudor evolved through a small-scale duplication from MSH6 to PDS5. However, convergent acquisition remains an alternative explanation, through convergent evolution that has led to similar amino acid Tudor domains of MSH6 and PDS5. **Second** (Figure 3, Step 2), the duplication of PDS5 into homologs A and B likely occurred between 360–480 MYA, during the early diversification of vascular plants, as exemplified by the presence of A/B types in basal lineages such as Polypodiopsida (ferns). This event marked the first emergence of the duplicated PDS5 genes in plants, analogous to the A/B duplication seen in vertebrates (Figure 3, Fig. 3). **Third** (Step 3), PDS5C-type homologs, characterized by shorter PDS5 domains while retaining the Tudor domain, emerged around 320–310 MYA in gymnosperms (seed plants), with subsequent expansion in angiosperms, most likely via duplication of PDS5A (Fig. 3). This sequential pattern highlights a plant-specific functional diversification of PDS5, with the acquisition of the Tudor histone-reader domain preceding the evolutionary origins of specialized homologs (PDS5C/E) with novel roles in epigenomic regulation and RdDM pathways (Göbel et al., 2024; Martín-Merchán et al., 2024; Niu et al., 2021; Quiroz et al., 2024).

**Figure 3.**
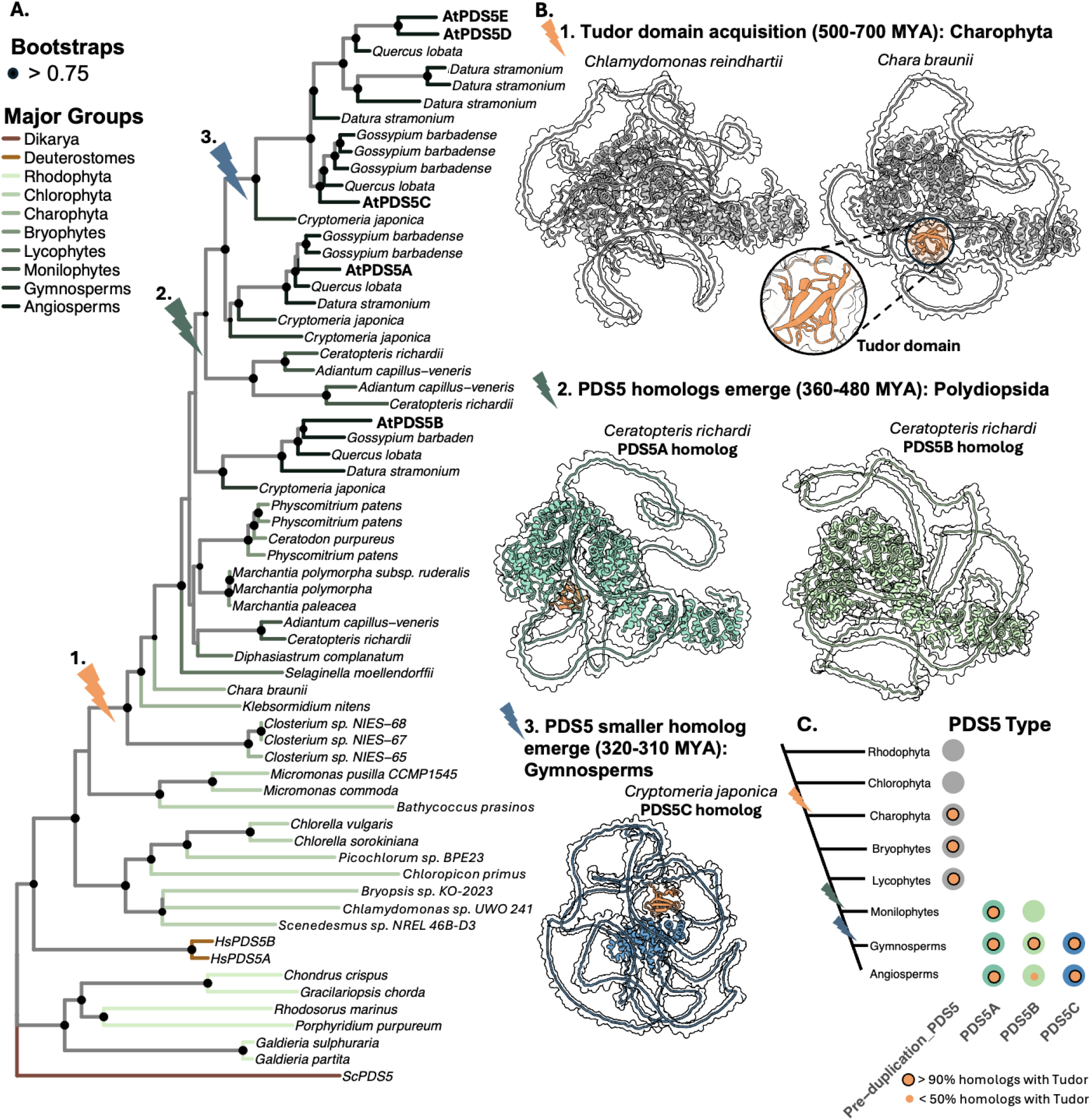
Maximum Likelihood Phylogenetic Tree of POSS Proteins Across Plantae. (A) Maximum-likelihood phylogenetic tree of the PDS5 protein family, constructed using the best-fit evolutionary model (*Q*.*insect*+*F*+*R20*) The tree is rooted in *Soccharomyces cerewwoe* PDS5 and includes 1–3 representative species per class. Bootstrap support values >0.75 (based on 1,000 replicates) are shown as black dots at the nodes. Major groups are color-coded in the branches. (B) Alphafold3 models of species with newly evolved features: 1. *Chlamydomonas reinhardtii* (Chlorophyta) and *Chara braunii* (Charophyta), with the Tudor domain structure highlighted in the inset; 2. *Ceratopteris richardii* (Monilophytes), showing PDS5A and PDS5B homologs; 3. *Cryptomeria japónica* (Gymnosperm), showing the PDS5C homolog. (C) Distribution of PDS5 types across clades. Circles represent the presence of Pre-duplication PDS5 (gray), PDS5A (teal), PDS5B (light green), and PDS5C (blue). An inner orange circle indicates the presence of the Tudor domain. Each circle denotes presence in at least one species per clade.

Notably, while the emergence of PDS5A and PDS5B in plants parallels the A/B duplication in vertebrates in evolutionary timing, sequence divergence patterns reveal a distinct contrast. Analyses of sequence conservation (Shannon entropy, Supplementary Fig. 6) show that plant PDS5 homologs, especially PDS5B, exhibit much higher sequence variability compared to their vertebrate counterparts. To rule out the possibility that these differences arose simply due to greater evolutionary divergence between the plant species sampled, we generated bootstrapped MSAs within key phyla (e.g., Streptophyta, Chordata, Ascomycota), sampling up to three species per class to maintain balanced taxonomic representation. These analyses confirmed that the sequence variability in plant PDS5B persists even when controlling for comparable evolutionary depths. Results represent the mean of 10 independent bootstrap replicates (Supplementary Fig. 6). This suggests that plant A/B homologs evolved with greater flexibility, potentially reflecting adaptation to plant-specific chromatin dynamics and regulatory processes.

### PDS5B Tudor Domains Lost H3K4me1-Affinity Following Duplication in Vascular Plants

To investigate the functional conservation of Tudor domains in plant PDS5 homologs, we aligned the predicted Tudor sequences of PDS5A, PDS5B, PDS5C, and pre-duplication Tudor domains. In Arabidopsis, the PDS5C Tudor domain was previously shown to bind H3K4me1 through an aromatic cage formed by three critical amino acids, facilitating cation-π interactions essential for recognition ((Niu et al., 2021); Fig. 4B). The importance of these amino acids has also been shown in the Tudor of MSH6, further validating the importance of these critical residues for specific histone binding (Monroe et al., 2025; Quiroz et al., 2024). Our analyses revealed that Tudor basal to the PDS5 gene duplication (pre-duplication Tudors) retain these key residues (Fig. 4A), supporting the idea that an early affinity for H3K4me was already established.

**Figure 4.**
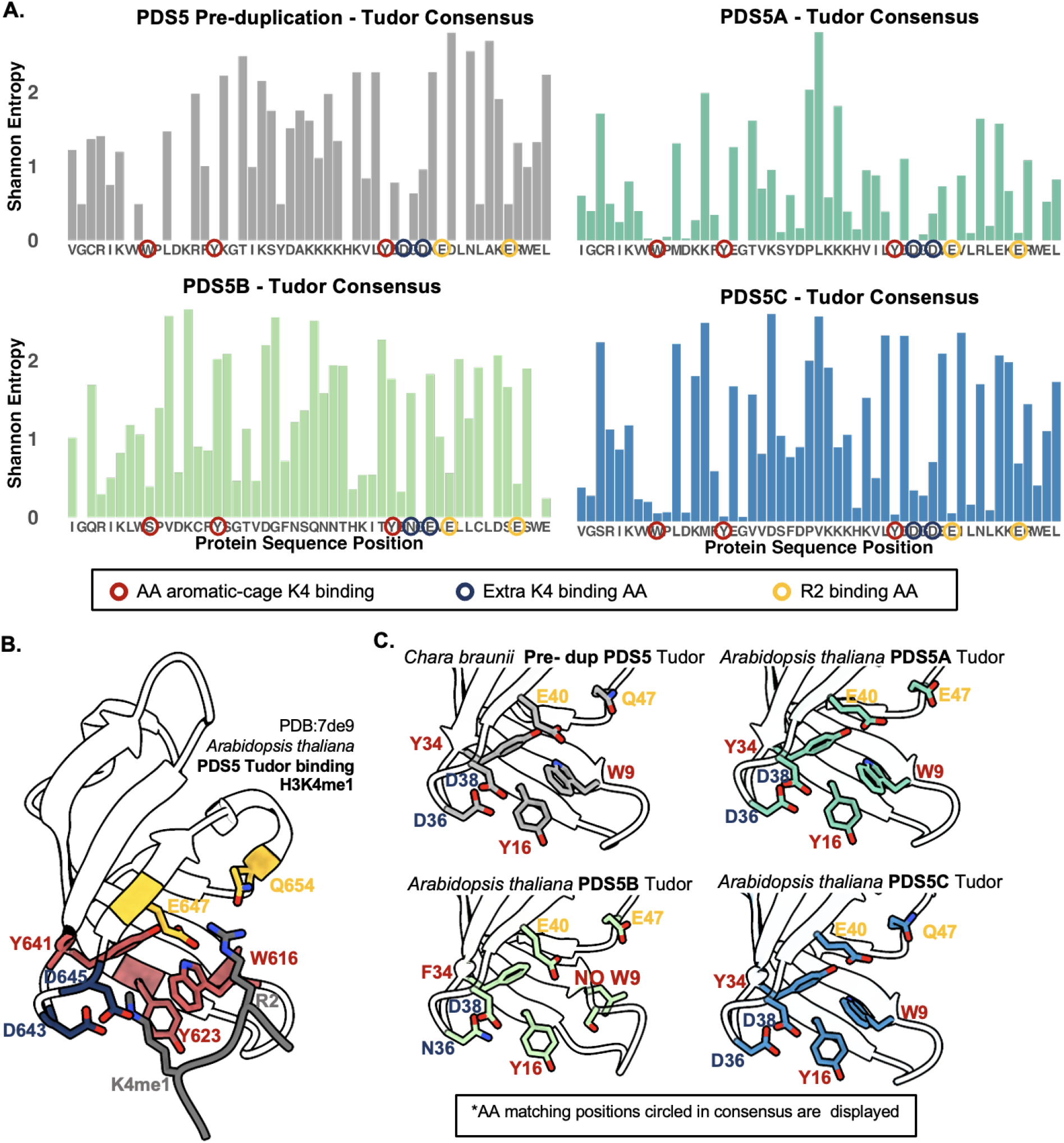
Evolutionary Analysis of the Tudor Domain in PDS5 Proteins Across Ptantae. (A) Shannon entropy plots for Tudor domains across PDS5 homologs (Pre-duplication, PDS5A, PDS5B and PDS5C). The consensus sequence without gaps for each homofog is displayed on the X-axis, and positions with functional importance are highlighted: aromatic cage residues for K4 binding (red circle), auxiliary K4 binding residues (blue circle), and R2 binding residues (yellow circle). (B) Structural model of the *Arabidopsis thaliana* PDS5C Tudor domain bound to H3K4mel (PDB: 7de9 (Niu et al. 2021)). highlighting key residues involved in Interaction with Histone K4mel Aromatic cage (Y641, W616, Y623) and auxiliary interactions (0&43, QMS), and key residues involved in Histone R2 interaction (E647, Q6S4). (C) Comparative structural analysis of the Tudor domains across clades, showing representative models from *Chara braunii* (pre-duplication), *Arabidopsis thaliana* POSSA, PDS5B, and PDS5C homologs Key residues matching portions in the consensus sequence (circled in (A) are visualized.

However, after the duplication event, divergence between Tudor domains of PDS5A and PDS5B homologs became evident. Among 432 total PDS5A homologs examined, 394 retained a predicted Tudor domain, with conservation of the key aromatic residues necessary for H3K4me1 binding. This is consistent with the adaptive value of PDS5 H3K4me1 interactions, at least for PDS5A and C homologs. In contrast, only 126 of 297 PDS5B homologs had a predicted Tudor domain. When present, Tudor domains lacked, particularly the residue corresponding to W9 (tryptophan), which is essential for H3K4me1 binding (Fig. 5A) (Niu et al., 2021; Quiroz et al., 2024). In these sequences, W9 was consistently replaced by a serine (S), indicating a loss of aromatic residue at this key site. This loss of affinity may help explain the complete loss of the Tudor domain in many of the predicted PDS5B homologs. However, PDS5B remains conserved across vascular plants, indicating adaptive retention of this paralog, potentially reflecting subfunctionalization or changes in subcellular localization. In summary, PDS5B is distinct from other PDS5 homologs in plants, showing evidence of relaxed constraint on an otherwise functional Tudor histone reader.

**Figure 5.**
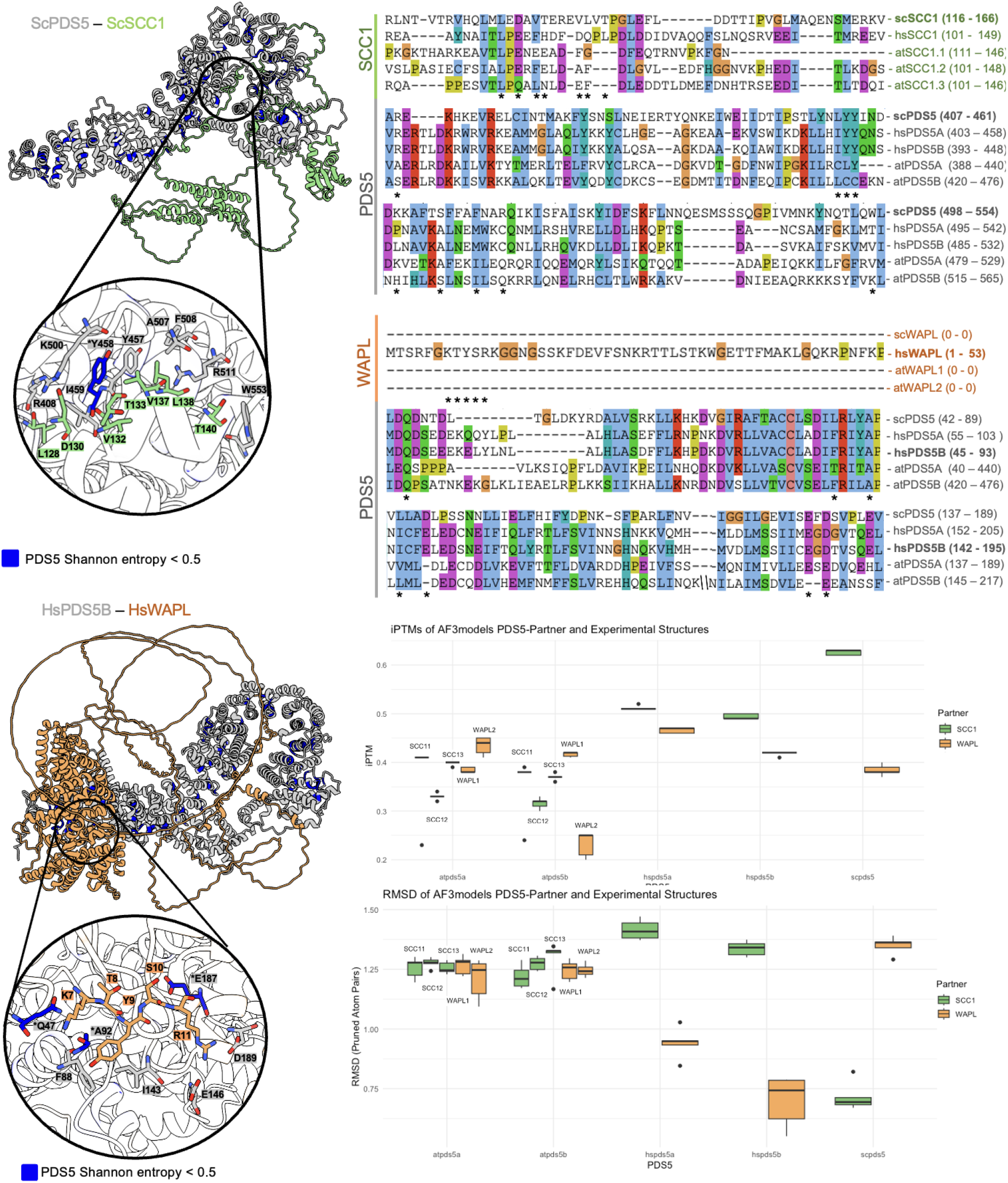
Structural and evolutionary comparison of the interaction between PDS5 and its partner proteins in Saccharomyces cerevisiae and Homo sapiens. (A) Structural models of **ScPDS5** - **ScSCC1** (in green) and **HsPDS5B** - **HsWAPL** (in orange) interactions, highlighting conserved regions in PDS5 proteins through Shannon entropy analysis (blue indicates low variability regions). A zoomed-in view of key interaction sites is shown for each complex. (B) Multiple sequence alignments of **POSS** and its partner proteins **(SCC1** in green and **WAPL** in orange), illustrating conservation and divergence of key residues marked with “ * “. (C) Stability assessment of POSS-partner interaction models through **pTM scores**, and structural similarity to experimental structures analyzed via **RMSD**. The interactions of PDS5 with SCC1 and WAPL are compared across species, revealing differences in interface flexibility and conservation.

### Structural Predictions Suggest Divergent Binding Modes of PDS5 With Cohesin Partners

We summarized known PDS5 interactors across humans, yeast, and Arabidopsis (Supplementary Table 1). Only SCC1 and WAPL have experimentally confirmed crystal structures with PDS5 (marked * in Supplementary Table 1); other interactions are supported mainly by indirect experimental evidence, such as co-immunoprecipitation. Notably, Sororin, an important cohesin regulator via its WAPL-antagonizing role (Lafont et al., 2010), is absent in *Saccharomyces cerevisiae* but was recently identified other fungi species and in Arabidopsis as AtSORORIN, expanding the recognized evolutionary conservation of this regulatory network (Prusén Mota et al., 2024). Nevertheless, experimental evidence directly linking PDS5 to many putative interactors in plants, such as SCC1, WAPL, and cohesin-associated factors, remains lacking. A key divergence remains the presence of PDS5C, a plant-specific homolog absent from animals and fungi, which links cohesin to RdDM pathways (Niu et al., 2021; Takei et al., 2024), highlighting lineage-specific innovations alongside conserved features across eukaryotes.

To explore the conservation and potential divergence of interaction interfaces, we focused on SCC1 and WAPL because they are the only PDS5 interactors with available experimental crystal structures: the yeast SCC1–PDS5 complex (PDB: 5F0N, 5F0O; Lee et al., 2016) and the human WAPL–PDS5 interaction (PDB: 5HDT; Ouyang et al., 2016). We compared these predicted binding regions across species via multiple sequence alignments(Fig. 5). While some SCC1-contact residues appear broadly conserved, many other contact points seem lineage-specific. This pattern may suggest that although the overarching PDS5–SCC1 binding mode is maintained, finer-scale interactions could vary across eukaryotes. For WAPL, while previous structural and biochemical studies (Ouyang et al., 2016) have identified a conserved PDS5-binding motif (the YSR motif) in vertebrates (critical for binding and resolved structurally), our comparative analysis reveals that this motif is absent in budding yeast and Arabidopsis WAPL homologs. Moreover, many amino acids essential for this interaction are poorly conserved in non-vertebrate PDS5 proteins. Interestingly, although the yeast WAPL-PDS5 complex has been shown to form and functionally counteract cohesion establishment (Sutani et al., 2009), the mode of interaction appears divergent, suggesting distinct binding interfaces evolved in fungi and plants.

We then used AlphaFold3 (Abramson et al., 2024) to model the ScPDS5-ScSCC1 (yeast), HsPDS5B-HsWAPL (human), and Arabidopsis complexes (Fig. 5A, C). We note that AlphaFold3 remains a hypothesis-generating tool, and experimental validation is required to confirm the precise nature of these interactions. We analyzed predicted Template Modeling (ipTM) scores, which estimate interface prediction confidence (range: 0–1; higher values indicate stronger confidence; Abramson et al., 2024). SCC1–PDS5 complexes in yeast and human yielded high ipTM scores (∼0.85–0.9), whereas WAPL–PDS5 models showed slightly lower values (∼0.65–0.75), consistent with previous findings that WAPL–PDS5 interactions are more transient or less stable (Kueng et al., 2006). In Arabidopsis, this trend was less pronounced: SCC1-PDS5 complexes exhibited overall lower ipTM scores, and WAPL2 appeared to show relatively higher predicted affinity for AtPDS5A, suggesting possible differences in interaction stability. RMSD analyses compared to known crystal structures further supported the predictive power of AlphaFold3 for yeast and human complexes, while higher RMSD values in Arabidopsis likely reflect genuine evolutionary divergence or reduced prediction accuracy due to the absence of plant-specific experimental structures. Although this accuracy may partly reflect training bias toward known structures, the distinct predicted interfaces across SCC1 and WAPL homologs suggest true evolutionary divergence. If AlphaFold3 were primarily template-driven, we might expect structural convergence; the observed variation might be reflecting real shifts in cohesin-PDS5 interaction modes across eukaryotes.

Altogether, these findings suggest a scenario in which PDS5 protein-protein interactions have evolved with divergent features across lineages. Our observations provide a starting point for future experiments to better understand the evolution and function of cohesin regulation.

## Conclusions

PDS5, known as the “cohesin gatekeeper,” is a highly conserved protein across the eukaryotic Tree of Life, confirming its indispensable role in regulating chromatin architecture and genome stability. Our findings reveal that while PDS5 is maintained, its evolutionary trajectory has diverged markedly across major eukaryotic lineages, reflecting lineage-specific adaptations.

In animals, the duplication of PDS5 into homologs A and B occurred early in vertebrate evolution. These two homologs have remained highly conserved, as shown by their low sequence divergence across vertebrate species. These findings support experimental evidence in mice, Xenopus, and humans that has demonstrated distinct yet complementary roles, suggesting subfunctionalization or neofuncitonalization while preserving critical cohesin-regulatory functions.

In contrast, plants exhibit a more dynamic evolutionary history. We identified the early acquisition of a Tudor histone reader domain in the ancestor of all land plants. This innovation likely linked PDS5 to histone-based epigenomic regulation, particularly through recognition of the H3K4me1 chromatin mark. This was followed by the emergence of duplicated homologs A and B during the diversification of vascular plants, and later, the evolution of specialized PDS5C-type homologs in seed plants, characterized by truncated PDS5 domains but retention of the Tudor reader. Notably, while vertebrate homologs have remained highly conserved, plant PDS5 homologs, especially PDS5B, display greater PDS5 sequence divergence. Interestingly, PDS5B has a degraded Tudor domain with indication of lost H3K4me1 affinity. Together, this evidence suggest a distinct adaptation of their cohesin-regulatory roles, this contrast is particularly striking given the comparable evolutionary timescales of homolog diversification in both lineages.

Our phylogenetic and structural analyses suggest that these divergent trajectories have led to evolving mechanisms of cohesin regulation, with modifications in protein-protein interaction interfaces and potential shifts in chromatin-binding dynamics. The emergence of distinct PDS5 homologs in both plants and vertebrates may represent a case of convergent evolution, enabling the fine-tuning of cohesin activity across different genomic contexts.

Beyond its classical role in cohesin regulation, the plant-specific PDS5C homolog exemplifies gene neofunctionalization: it participates in the RNA-directed DNA methylation (RdDM) pathway, bridging cohesin dynamics with epigenomic silencing mechanisms. This dual function highlights the evolutionary plasticity of PDS5, allowing it to integrate cohesin-related processes with lineage-specific regulatory networks.

Together, these findings reinforce PDS5 as a model of both evolutionary conservation and innovation, offering new insights into how essential chromatin regulators adapt and diversify across eukaryotes.

## Materials and Methods

### Putative Homolog Identification and Filtering

PDS5 homologs were identified across eukaryotic proteomes using BLASTp (Protein-Protein BLAST 2.15.0) with yeast, human, and Arabidopsis PDS5 sequences as queries (Uniprot accession: Q04264, Q29RF7, Q9NTI5, B6EUB3, F4I735, Q8GUP3, A8MRD9, Q9S9P0). Searches were performed against the NCBI eukaryotic protein database, and hits with E-values below 1e-6 were retained. To reduce redundancy, sequences within each TaxID were aligned using Clustal Omega (version 1.2.4), and proteins sharing >85% sequence identity were clustered, retaining the longest representative per cluster, thereby minimizing the inclusion of truncated or incomplete protein versions (Supplementary Fig. 1A).

### Taxonomic Classification

We first obtained genome-to-species associations using NCBI genome annotation files, linking analyzed PDS5 homologs to their respective species and TaxIDs. Next, we used the taxonomizr R package (version 0.10.6) to retrieve full taxonomic classifications (phylum, class, order, family, genus, species) for each TaxID. Where taxonomic information was missing or incomplete, we performed additional curation by manually reviewing and resolving overlapping codes and harmonizing inconsistent annotations. This finalized taxonomy table provided the framework for grouping sequences into major eukaryotic lineages (Protists, Fungi, Animalia, Plantae) and guided the subsequent subsampling and phylogenetic analyses.

### Domain Prediction and Filtering

Domain composition was analyzed using InterProScan (version 5.72-103.0), retaining sequences containing the PDS5 domain (Pfam PF20168) with E-values < 0.1. Tudor (CDD cd20404) and IDR (MobiDBLite) were annotated when present. To exclude likely truncated annotations, we applied group-specific domain size thresholds based on the distribution of PF20168 sizes per major eukaryotic clade (Supplementary Fig. 1B). For Protists and Fungi, sequences >800 amino acids were retained; for Animals, sequences between 900 and 1,200 amino acids; and for Plantae, sequences between 900-1,200 and 150-300 amino acids, reflecting the dual distribution observed in plant PDS5 homologs. These thresholds prioritized conserved features and minimized potential annotation artifacts.

### Phylogenetic Analysis

To infer the evolutionary relationships of PDS5 homologs, we performed multiple sequence alignments (MSA) of curated PDS5 domain sequences (Pfam PF20168) using Clustal Omega (version 1.2.4) with default parameters. To ensure balanced representation across major eukaryotic lineages, we applied a subsampling strategy: for each major group, up to three representative species per taxonomic class were selected, prioritizing taxonomic diversity and well-annotated genomes. This was implemented using a custom Python script, which bootstrapped three independent datasets to capture diversity while maintaining manageable tree sizes.

Maximum likelihood phylogenetic trees were constructed using IQ-TREE multicore version 2.4.0, with the best-fit substitution model automatically selected under the Bayesian Information Criterion (BIC). Node support was assessed using ultrafast bootstrap approximations with 1000 replicates to ensure statistical robustness. Trees were visualized and annotated using the R package ggtree (version 3.6.2), with major eukaryotic clades (Protists, Fungi, Animalia, Plantae) color-coded for clarity. Basal lineages were highlighted to emphasize early-diverging groups, and representative species were labeled at the tips. The final phylogenetic tree presented in Fig. 2A reflects the evolutionary clustering of PDS5 homologs across a wide spectrum of eukaryotic diversity.

### Comparative Analyses of PDS5 Homologs

Shannon entropy analysis was performed on multiple sequence alignments (MSAs) of PDS5 domains from homologs in plants and animals to assess the conservation and variability of residues across taxa. Separate alignments were conducted for plant and animal homologs, focusing on conserved structural and functional regions of the PDS5 domain. For the PDS5 domain specifically, bootstrapped alignments were generated from a curated dataset representing key taxonomic groups (e.g., Ascomycota, Chordata, and Streptophyta). To create multiple bootstrap replicates, a Python pipeline was employed to sample sequences within each phylum, grouping by taxonomic class and including reference sequences. These FASTA files were then aligned using Clustal Omega, producing both Clustal and FASTA format alignments.

Subsequently, custom R scripts were used to calculate Shannon entropy on a per-column basis by computing the amino acid frequency distributions (excluding gaps).

For plants, additional MSAs were created for homologs with annotated Tudor domains. We classified plant PDS5 homologs into pre-duplication, A, B, and C-type groups. Tudor domains from homologs preceding the A/B duplication event were classified as pre-duplication Tudor based on taxonomic inferences. While A, B, and C-type were classified with a custom HMMR. This model was built using PDS5 homologs sequences from five representative angiosperm species. MSAs were generated for each Homolog-Tudor group using Clustal Omega, and Shannon entropy and amino acid frequency distributions were calculated for each.

### Protein-Protein Interaction Prediction

Structural models of PDS5-partner protein complexes were generated using AlphaFold3 (Abramson et al., 2024). We focused on two well-characterized interactions: the yeast ScPDS5-ScSCC1 complex and the human HsPDS5B-HsWAPL complex, as these have available experimental structures for benchmarking. Models were built for the full-length proteins and their interactors to predict binding interfaces and complex stability.

To assess conservation across lineages, Shannon entropy per amino acid was calculated from multiple sequence alignments (MSAs) of PDS5 homologs across three major phylogenetic groups: Ascomycota (yeast), Streptophyta (plants), and Chordata (vertebrates). These entropy scores were overlaid onto the predicted structures to highlight conserved vs. variable regions within the binding interfaces.

Interaction model confidence was evaluated using predicted Template Modeling (pTM) scores, which estimate the reliability of inter-chain contacts within the complex (range 0–1, with higher values indicating greater confidence). Structural accuracy was assessed via root-mean-square deviation (RMSD), calculated using the MatchMaker tool in ChimeraX, by aligning AlphaFold3-predicted complexes against available experimental structures: PDB 5H68 (ScPDS5–ScSCC1; Lee et al., 2016) and PDB 6H8Q (HsPDS5B–HsWAPL; Ouyang et al., 2016).

Finally, interaction surfaces were compared across species through MSAs of key binding regions, using UniProt-accessioned sequences of SCC1, WAPL, and PDS5 homologs in humans, yeast, and Arabidopsis. The aim was to identify conserved vs. lineage-specific contact residues that might underlie evolutionary divergence in cohesin regulation.

### Visualization

Phylogenetic and entropy analyses were visualized using R packages, including ggtree for phylogenetic trees and ggplot2 for entropy distributions. Protein interaction models were visualized in ChimeraX, highlighting key interaction residues and domain alignments. This comprehensive pipeline enabled the identification, functional annotation, and evolutionary analysis of PDS5 proteins across eukaryotes, providing insights into their structural conservation and functional diversification. Further details, including E-values, database versions, and parameter settings, are provided in the supplementary information.

## Acknowledgments

This work was supported by NSF Grants 2317191 and 2338236 to J.G.M. and by Beca Chile doctoral fellowship (ANID) to D.Q Research was conducted at the University of California Davis, which is located on land that was the home of the Patwin people for thousands of years.

## Code availability

Code for this research is maintained at https://github.com/DaniPQ/PDS5_Evolution This repository contains all the scripts, intermediate tables, and summary datasets used in the evolutionary analysis of PDS5 proteins

## Supplemental Materials

### Supplemental materials for

**Title**: Evolutionary trajectories of cohesin gatekeepers across the Tree of Life

**Authors**: Daniela Quiroz^1^, Alice Pierce^1^, Grey Monroe^1^

## Supplemental Datasets

The following supplementary datasets are provided in the GitHub repository DaniPQ/PDS5_Evolution:

**Supplementary Dataset 1:** Contains raw and merged BLASTp + HMMER homolog identification tables, including non-redundant TaxID annotations across eukaryotic proteomes.

**Supplementary Dataset 2:**Provides final curated species-level tables and taxonomic databases used to guide phylogenetic tree construction and downstream analyses.

**Supplementary Dataset 3:**Includes InterProScan domain prediction outputs and size-filtered results, focusing on PDS5 and Tudor domain identification.

**Supplementary Dataset 4:** Contains Shannon entropy and consensus sequence tables for specific PDS5-types (Preduplication, PDS5A, PDS5B, and PDS5C) Tudor domains.

**Supplementary Dataset 5:** Summarizes detailed entropy results for PDS5 homologs across major clades (yeast, fungi, plants, animals), highlighting post-duplication evolutionary patterns.

**Supplementary Dataset 6:** Contains AlphaFold3-based structural comparison results, including RMSD and IPTM metrics for PDS5 interactions with SCC1, WAPL, and related partners.

## Supplemental figures

**Supplementary Figure 1.**
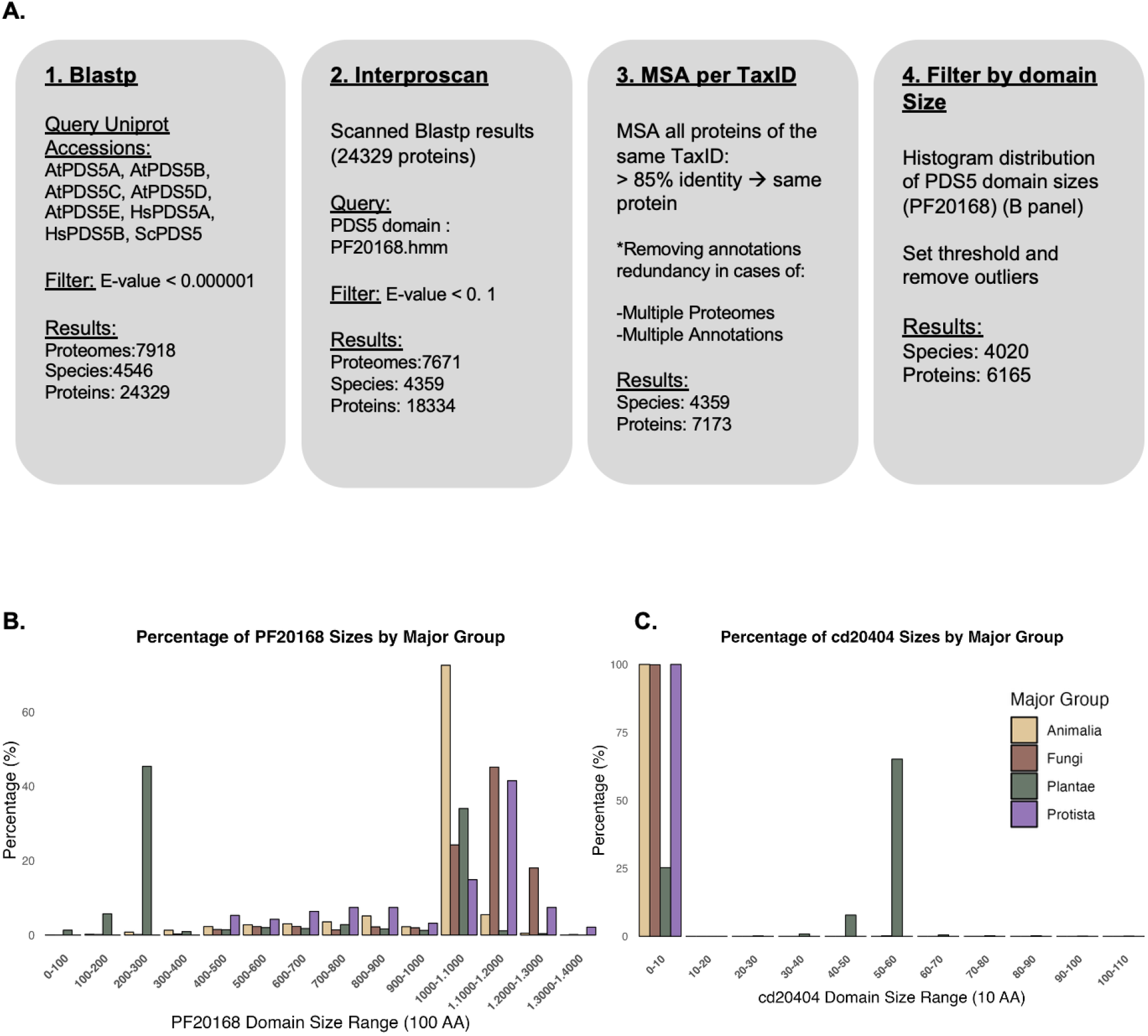
Pipeline for putative PDS5 homologs identification. (A) Overview of the computational workflow for identifying PDS5 homologs across eukaryotic proteomes The process involves four main steps: 1. Blastp, querying Uniprot accessions (AtPDS5A, AtPDS5B. AtPDS5C, AtPDS5D, AtPDS5E, HsPDS5A, HsPDS5B, ScPDS5) against 8,538 eukaryotic proteomes, filtered by an E-value < 0.000001, yielding 7,918 proteomes, 4,546 species, and 24,329 proteins. 2. InterProScan, scanning Blastp hits for the PF20168 domain with an E-value threshold of < 0.1, resisting in 7,671 proteomes, 4,359 species, and 18,334 proteins. 3. MSA per TaxID, grouping all proteins within the same TaxlD with >85% identity Into single proteins, removing redundancy from multiple annotations or proteomes, yielding 4,359 species and 7,173 proteins. 4. Doman Size Filtering, producing histograms for PF20168 and cd20404 domain see distributions, resulting in 4,020 species and 6,165 proteins after setting thresholds and removing outliers (possible truncated predictions! (B) Percentage distribution of PF20168 domain sees (100 AA bins) across major groups (Protista, Fungi, Animalia, Plantae). (C) Percentage distribution of cd20404 domain sizes (10 AA bins) across major groups.

**Supplementary Figure 2.**
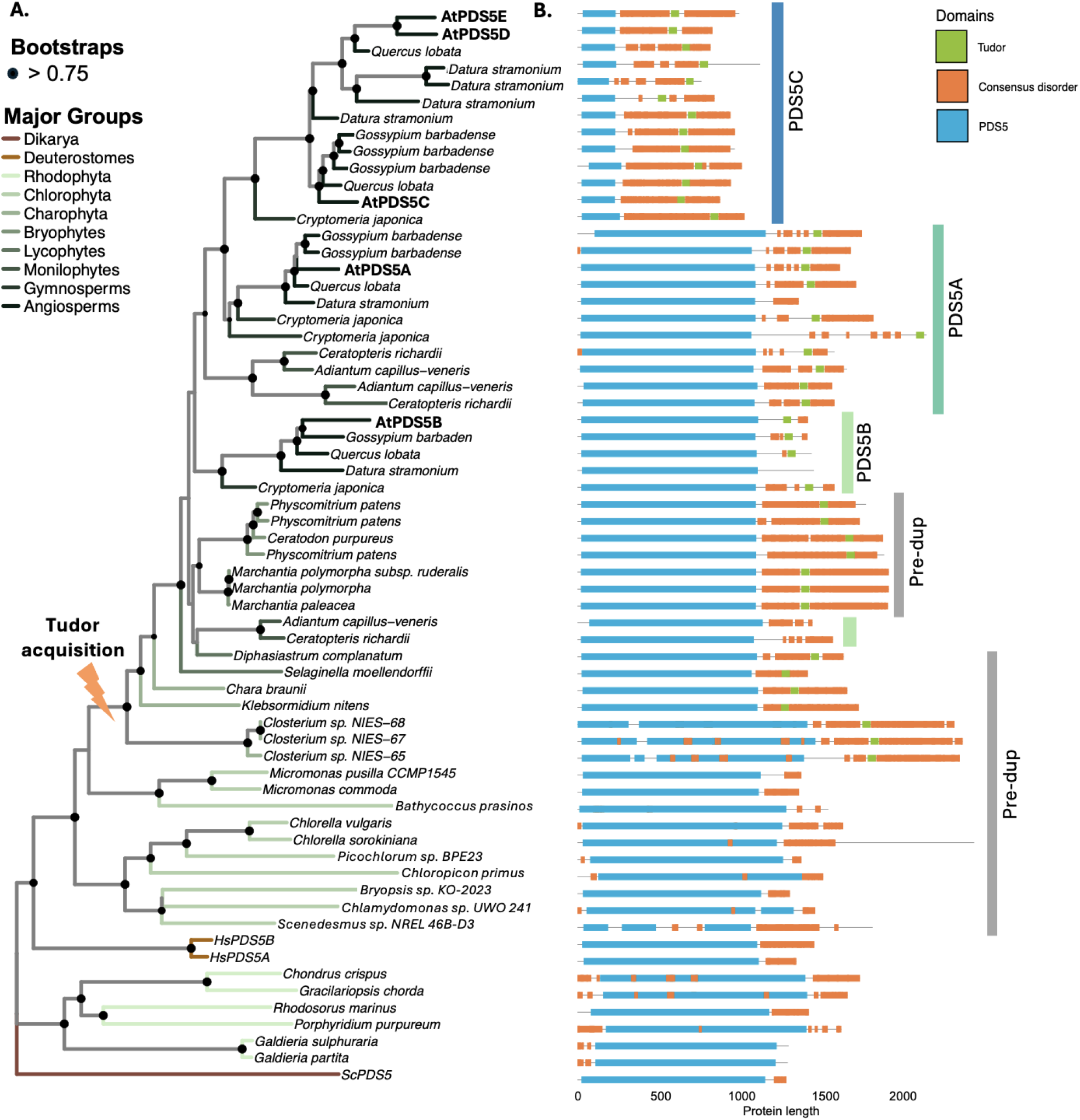
Maximum Likelihood Phylogenetic Tree of PDS5 Proteins Across Plantae. **(A)** Maximum-likelihood phylogenetic tree of the PDS5 protein family, constructed using the best-fit evolutionary model *(Q*.*insect+F+R10)*. The tree is rooted in *Saccharomyces cerevisiae* PDS5 and includes 1–3 representative species per class. Bootstrap support values >0.75 (based on 1,000 replicates) are shown as black dots at the nodes. Major groups are color-coded in the branches. **(B)** Domain architecture of PDS5 proteins based on lnterProScan results. The domains are highlighted: Tudor domains (green), consensus disordered regions (orange) and PDS5 domains (blue). Positions correspond to sequence coordinates from 1 to 2000 amino acids.

**Supplementary Figure 3.**
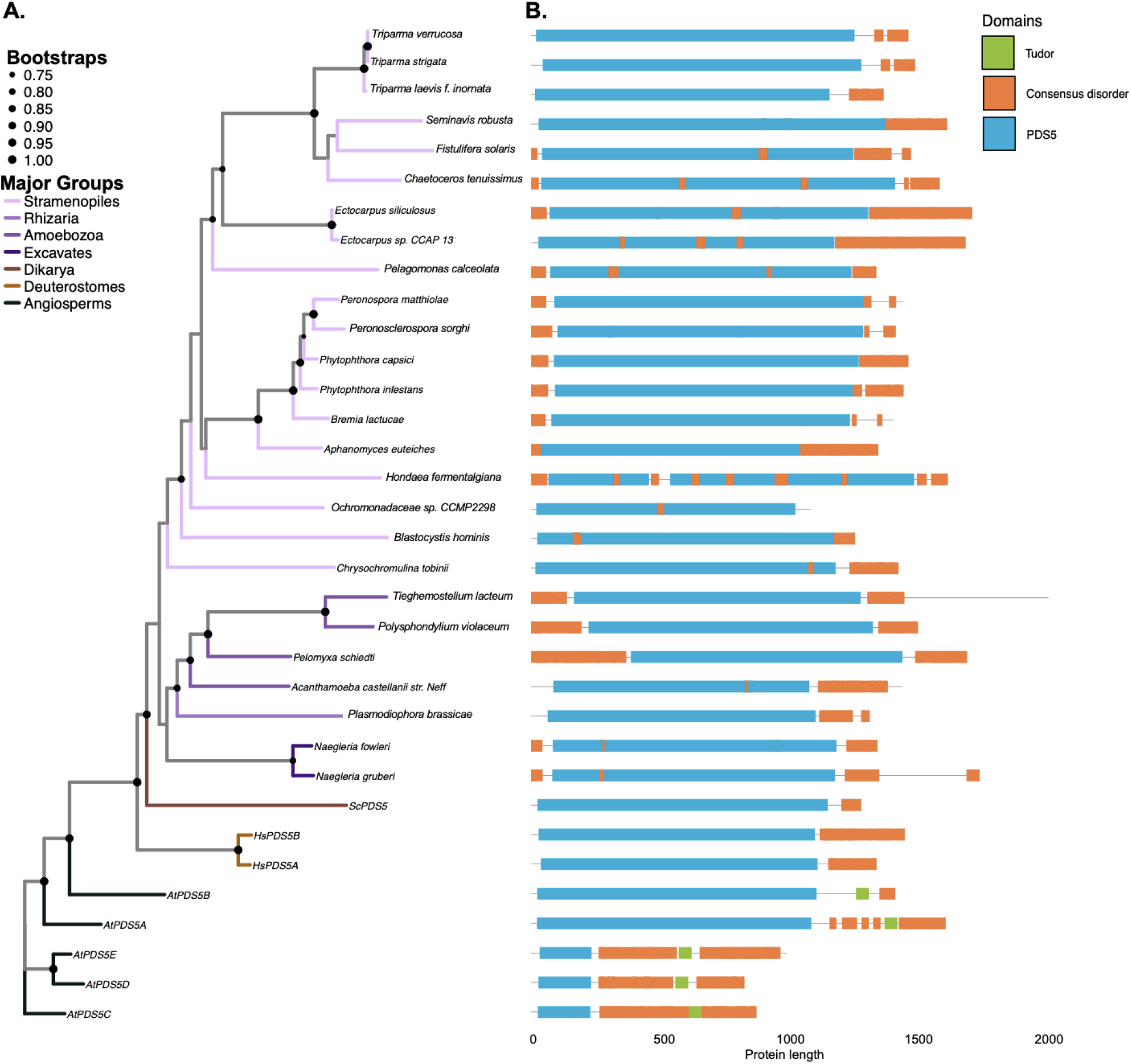
Maximum Likelihood Phylogenetic Tree of POSS Proteins Across Protists. (A) Maximum-likelihood phylogenetic tree of the POSS protein family, constructed using the best-fit evolutionary model *(Q*.*insect+F+RlO)*. The tree is rooted in *Arabidopsis tholiano* POSS and includes 1–3 representative species per class. Bootstrap support values >0.75 (based on 1,000 replicates) are shown as black dots at the nodes. Major groups are color-coded in the branches. (B) Domain architecture of POSS proteins based on lnterProScan results. The domains are highlighted: Tudor domains (green), consensus disordered regions (orange) and POSS domains (blue). Positions correspond to sequence coordinates from 1 to 2000 amino acids.

**Supplementary Figure 4.**
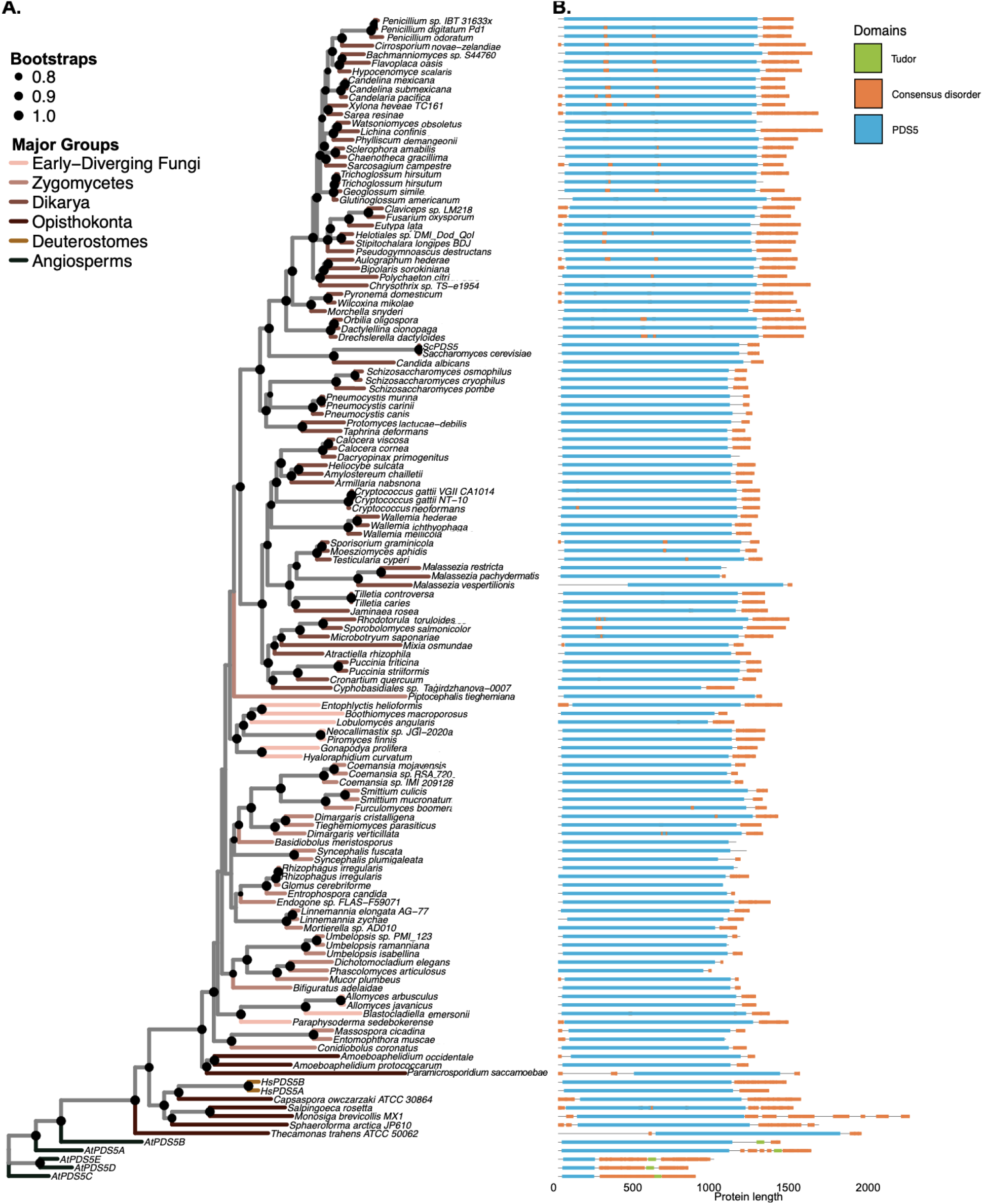
Maximum Likelihood Phylogenetic Tree of POSS Proteins Across Fungi. (A) Maximum-likelihood phylogenetic tree of the PDS5 protein family, constructed using the best-fit evolutionary model *(Q*.*insect+F+R10)*. The tree is rooted in *Arabidopsis tha/iana* PDS5 and includes 1–3 representative species per class. Bootstrap support values >0.75 (based on 1,000 replicates) are shown as black dots at the nodes. Major groups are color-coded in the branches. (B) Domain architecture of PDS5 proteins based on lnterProScan results. The domains are highlighted: Tudor domains (green), consensus disordered regions (orange) and PDS5 domains (blue). Positions correspond to sequence coordinates from 1 to 2000 amino acids.

**Supplementary Figure 5.**
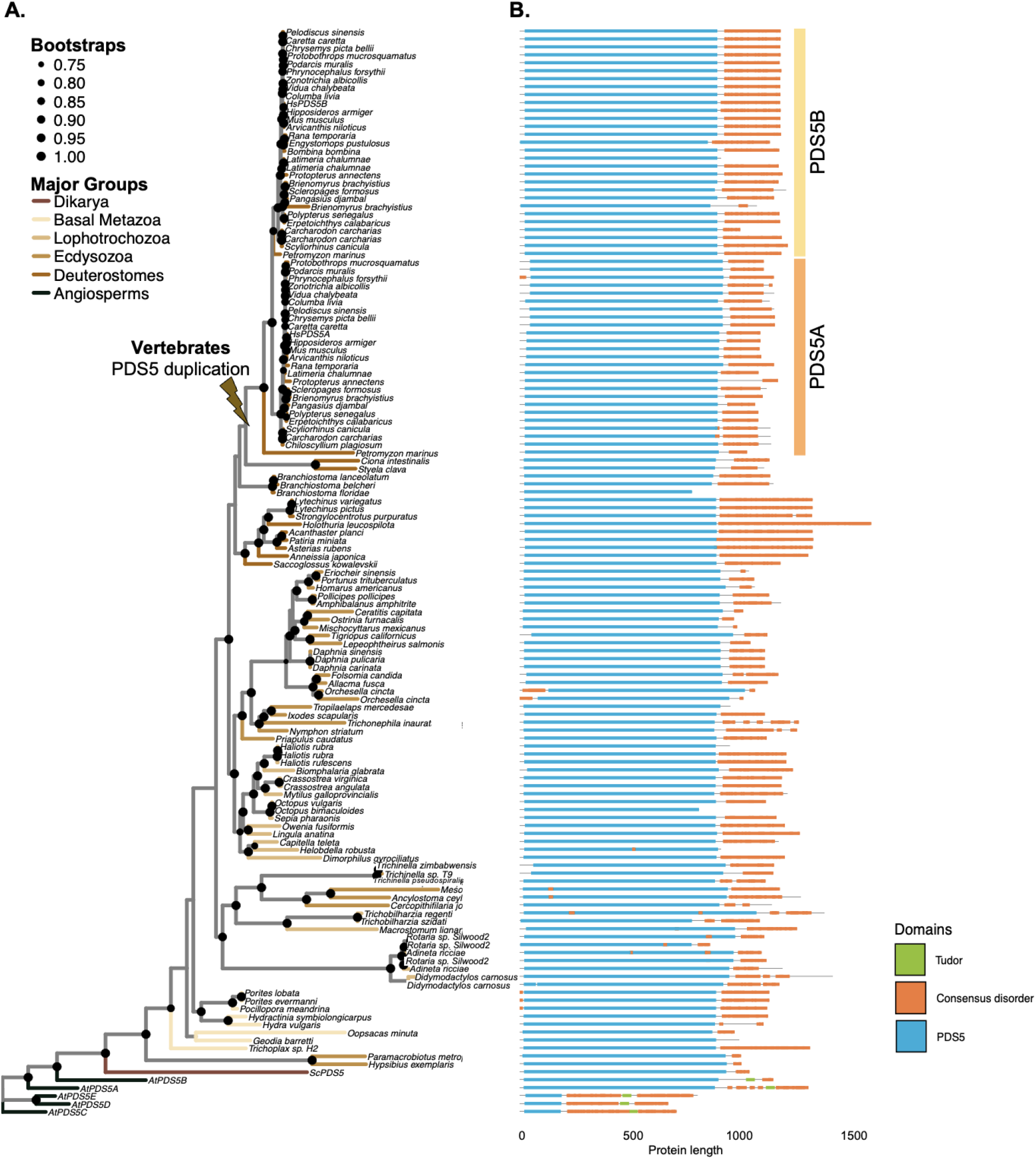
Maximum Likelihood Phylogenetic Tree of PDS5 Proteins Across Animalia. **(A)** Maximum-likelihood phylogenetic tree of the POSS protein family, constructed using the best-fit evolutionary model *(Q*.*insect+F+R10)*. The tree is rooted in *Arabidopsis thaliana* POSS and includes 1–3 representative species per class. Bootstrap support values >0.75 (based on 1,000 replicates) are shown as black dots at the nodes. Major groups are color-coded in the branches. (B) Domain architecture of POSS proteins based on lnterProScan results. The domains are highlighted: Tudor domains (green), consensus disordered regions (orange) and POSS domains (blue). Positions correspond to sequence coordinates from 1 to 2000 amino acids.

**Supplementary Figure 6.**
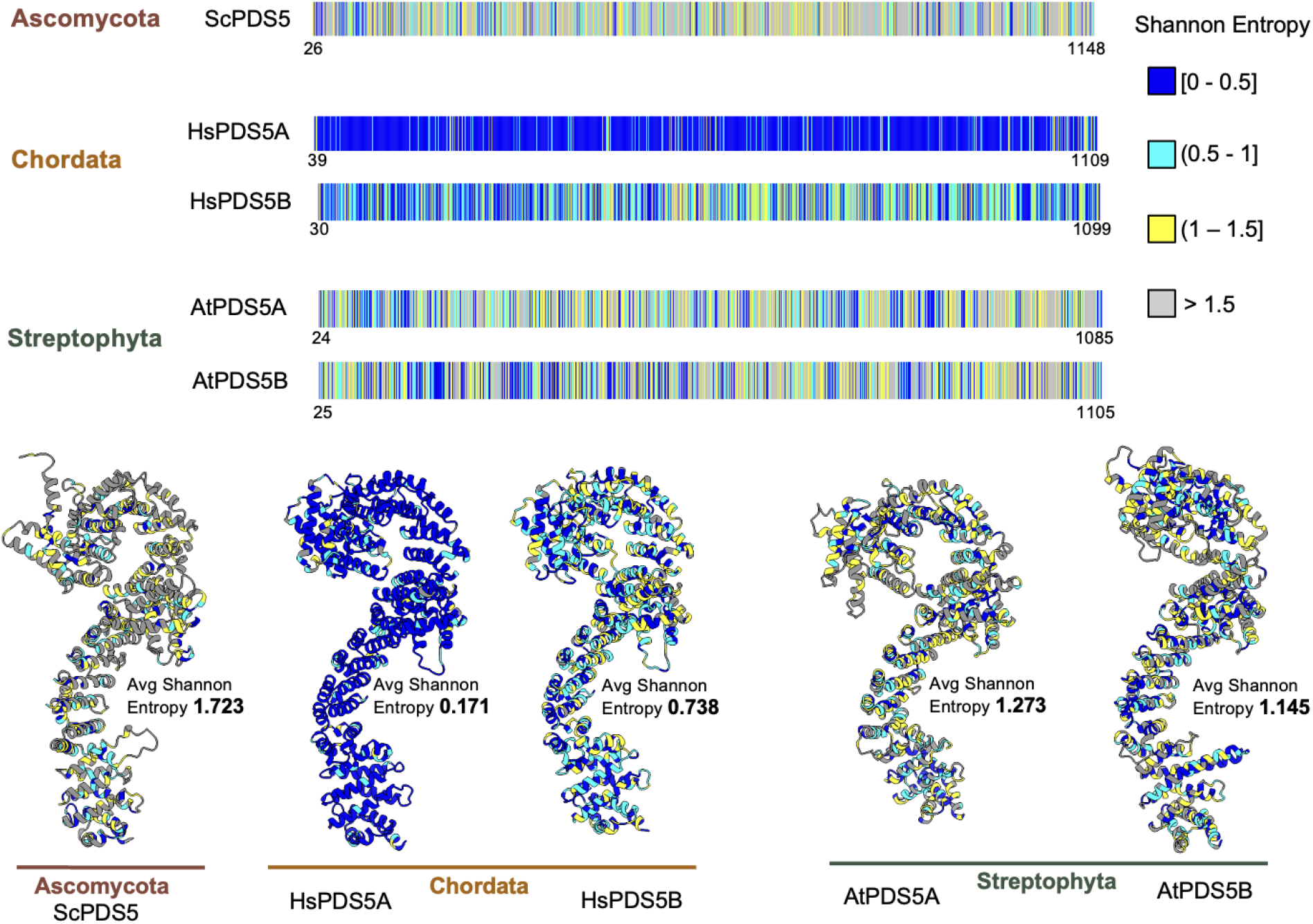
Shannon Entropy Distribution and 3D Structures of PDS5 Proteins. Top panels (horizontal colored strips) represent Mean Shannon Entropy per amino acid position across 10 bootstrapped alignments. Each rectangle corresponds to one alignment set: Ascomycota (ScPDS5), Chordata (HsPDS5A, HsPDS5B), Streptophyta (AtPDS5A, AtPDS5B). Colors indicate entropy ranges: blue [0, 0.5), cyan (0.5, 1), yellow (1, 1.5), and grey > 1.5. Positions with higher entropy (>1.5) suggest greater sequence variability, while lower entropy (<0.5) indicates more conservation. Lower panels show the predicted 3D structures (AlphaFold3) for each PDS5 variant, colored according to the same four entropy intervals. The overlay of entropy onto the protein backbone highlights regions of high or low sequence conservation across each phylogenetic group.

## Supplemental tables

**Supplementary Table 1.** Comparative summary of known and predicted PDS5 protein interactors across Human. *Saccharomyces cerevisiae*, and *Arabidopsis thaliana*

|  | Homologs(Uniprot) |  |  | Interaction evidence |  |  |  |  | Biological Function |
| --- | --- | --- | --- | --- | --- | --- | --- | --- | --- |
|  |  |  |  | Human |  | Yeast | Arabidopsis |  |  |
|  | Human | Yeast | Arabidopsis | PDS5A | PDS5B | Pds5 | PDS5A | PDS5B |  |
| SMC1 | Q14683, Q8NDV3 | P32908 | Q6Q1P4 | Yes, No | Yes, No | Yes | No | No | Cohesin complex<br>SMC1B, meiosis specific |
| SMC3 | Q9UQE7 | P47037 | Q56YN8 | Yes | Yes | Yes | No | No | Cohesin complex |
| SCC1/<br>RAD21/<br>MCD1/<br>SYN | O60216 | Q12158 | Q9S7T7,<br>Q9FQ20,<br>Q9FQ19,<br>Q8W1Y0 | Yes | Yes | Yes* | No | No | Cohesin complex |
| SCC3/<br>STAG/<br>SA/<br>IRR1 | Q8WVM7,<br>Q8N3U4 | P40541 | O82265 | Yes, Yes | Yes, Yes | Yes | No | No | Cohesin complex |
| SCC4/<br>MAU2 | Q9Y6X3 | P40090 | Q9FGN7 | No | No | No | No | No | Cohesin complex |
| SCC2/<br>NIPBL | Q6KC79 | Q04002 | A5HEI1 | Yes | Yes | Yes | No | No | Cohesin complex |
| WAPL/<br>RAD61/<br>Wpl1 | Q7Z5K2 | Q99359 | F4I7C7,<br>Q9C951 | Yes | Yes* | Yes | No | No | Cohesin complex |
| ESCO/<br>ECO1/<br>CTF7 | Q5FWF5,<br>Q56NI9 | P43605 | A7UL74 | Yes | No | Yes | No | No | Cohesin complex |
| CDCA5/<br>Sororin | Q96FF9 | No | No | Yes | Yes | No | No | No | Cohesin complex |
| Separin/<br>ESP1/<br>RSW4/<br>Separase/<br>ESPL1 | Q14674 | Q03018 | Q5IBC5 | No | No | No | No | No | Cohesin complex |
| REC8 | O95072 | Q12188 | No | No | No | Yes | No | No | Cohesin complex (meiosis specific) |
| SMC5 | Q8IY18 | Q08204 | Q9LFS8 | No | No | Yes | No | No | SMC complex - DNA repair |
| SWAP1 | Q6NVH7 | No | No | No | Yes | No | No | No | ATPase involved in homologous recombination. |
| BRCA2/<br>BRC2 | P51587 | No | Q7Y1C5,<br>Q7Y1C4 | No | Yes | No | No | No | Involved in double-strand break repair |
| TOP2 | P11388,<br>Q02880 | P06786 | P30182 | No | No | Yes | No | No | DNA Topoisomerase 2 |
| DEK3 | P35659 | No | Q9SUA1 | No | No | No | Yes | No | Contributes to the modulation of chromatin structure |
| PDS5C | No | No | Q8GUP3 | No | No | No | Yes | No | PDS5 plant specific homolog - RdDM - Cohesin |

Uniprot codes shown. \*Indicates proteins with solved crystal structures in complex with PDS5 (Lee et al., 2016; Ouyang et al., 2016). Interaction data compiled from Protein Atlas, SGD, TAIR, STRING, IntAct, and recent literature (Lafont et al., 2010; Niu et al., 2021; Takei et al., 2024; Prusén Mota et al., 2024).

